# Polymorphic Structure Determination of the Macrocyclic Drug Paritaprevir by MicroED

**DOI:** 10.1101/2023.09.09.556999

**Authors:** G Bu, E Danelius, L.H Wieske, T Gonen

**Affiliations:** Department of Biological Chemistry, University of California Los Angeles, 615 Charles E.Young Drive South, Los Angeles, CA 90095, USA; Howard Hughes Medical Institute, University of California Los Angeles, Los Angeles, CA 90095, USA; Department of Chemistry – BMC, Uppsala University, Husargatan 3, 75237 Uppsala, Sweden; Department of Physiology, University of California Los Angeles, 615 Charles E. Young Drive South, Los Angeles, CA 90095, USA

**Keywords:** MicroED, CryoEM, Macrocycles, Polymorphism, Molecular Chameleons, HCV Protease

## Abstract

Paritaprevir is an orally bioavailable, macrocyclic drug used for treating chronic Hepatitis C virus infection. Its structures had been elusive to the public until recently when one of the crystal forms was solved by MicroED. In this work, we report the MicroED structures of two distinct polymorphic crystal forms of paritaprevir from the same experiment. The different polymorphs show conformational changes in the macrocyclic core, as well as the cyclopropylsulfonamide and methylpyrazinamide substituents. Molecular docking shows that one of the conformations fits well into the active site pocket of the NS3/4A serine protease target, and can interact with the pocket and catalytic triad via hydrophobic interactions and hydrogen bonds. These results can provide further insight for optimization of the binding of acylsulfonamide inhibitors to the NS3/4A serine protease. In addition, this also demonstrate the opportunity of deriving different polymorphs and distinct macrocycle conformations from the same experiments using MicroED.

## Introduction

Polymorphism, the phenomenon where a solid chemical compound occurs in more than one crystalline form with different molecular packing and/or conformation can result in different drug properties such as solubility, stability, bioavailability, potency, and toxicity, and is therefore of high importance to pharmaceutical research and development. [1-3] Active pharmaceutical ingredients (APIs) in pure drugs or formulated products are commonly available to patients as solid forms, thus it is of critical importance to understand and control polymorphism for optimal drug performance. [4, 5] Polymorphism can be characterized by a combination of techniques including X-ray diffraction (single crystal or powder) and spectroscopy (Raman, infrared or NMR), [2] and insufficient characterization occasionally can cause inactivation of drugs for life-threatening diseases. One infamous example is the anti-AIDS drug ritonavir which was commercialized based on the only previously described crystal form. The discovery of a polymorph in the commercial capsules with reduced solubility and bioavailability led to its temporary withdrawal from the market. [6] Molecular properties such as size and conformational flexibility can strongly influence the propensity of forming crystalline polymorphs, hence the elucidation of polymorphism is of higher importance outside the classical rule of five (Ro5) space for drug development.

Macrocycles are defined as having a cyclic core with 12 or more heteroatoms and represents novel chemical modalities beyond the traditional rule of 5 (bRo5) for oral absorption. [7, 8] Compared with classic small molecule drugs within Ro5, macrocycles are more flexible and complex, can adopt multiple conformations and bind to difficult-to-drug protein targets including those with high mutation rates or flat surfaces. [9, 10] Due to the conformational flexibility of macrocyclic drugs, they are prone to be crystalized as polymorphs. [5] Paritaprevir (ABT-450, Figure 1) is one of the highest molecular weight drugs approved for oral administration. [11] It was developed by AbbVie and Enanta Pharmaceuticals as a potent inhibitor of hepatitis C virus (HCV) non-structural 3/4A protease which plays a vital role in viral replication and assembly. [12, 13] Paritaprevir was approved as a component of direct-acting anti-HCV combination therapy under the brand names Viekira Pak (along with ombitasvir, ritonavir and dasabuvir) and Technivie (along with ombitasvir and ritonavir). [13, 14] Despite its wide use in clinic, until recently there were no published structures of paritaprevir in the Cambridge Crystallographic Data Center (CCDC) or target bound structures in the Protein Data Bank (PDB), illustrating the difficulty in studying theses large and flexible macrocyclic structures. Using a SerialEM-based high-throughput microcrystal electron diffraction (MicroED) data collection, we recently solved one of the crystal forms of paritaprevir, hereafter referred to as form α. [15] MicroED is a cryogenic electron microscopy (cryo-EM) method for structure determination from tiny crystals of micron to sub-micron size, bypassing the crystallization assay required for conventional X-ray diffraction. [16-18] Recent works has shown the ability of MicroED to solve polymorphic structures of small molecules including glycine by time-resolved in situ crystallization, [19] diketopyrrolopyrroles by drop casting on EM grids, [20] indomethacin by crystallization screening, [21] bis-arylacylhydrazone in the presence or absence of hydrates, [22] and vemurafenib by melt crystallization. [23]

**Figure 1.**
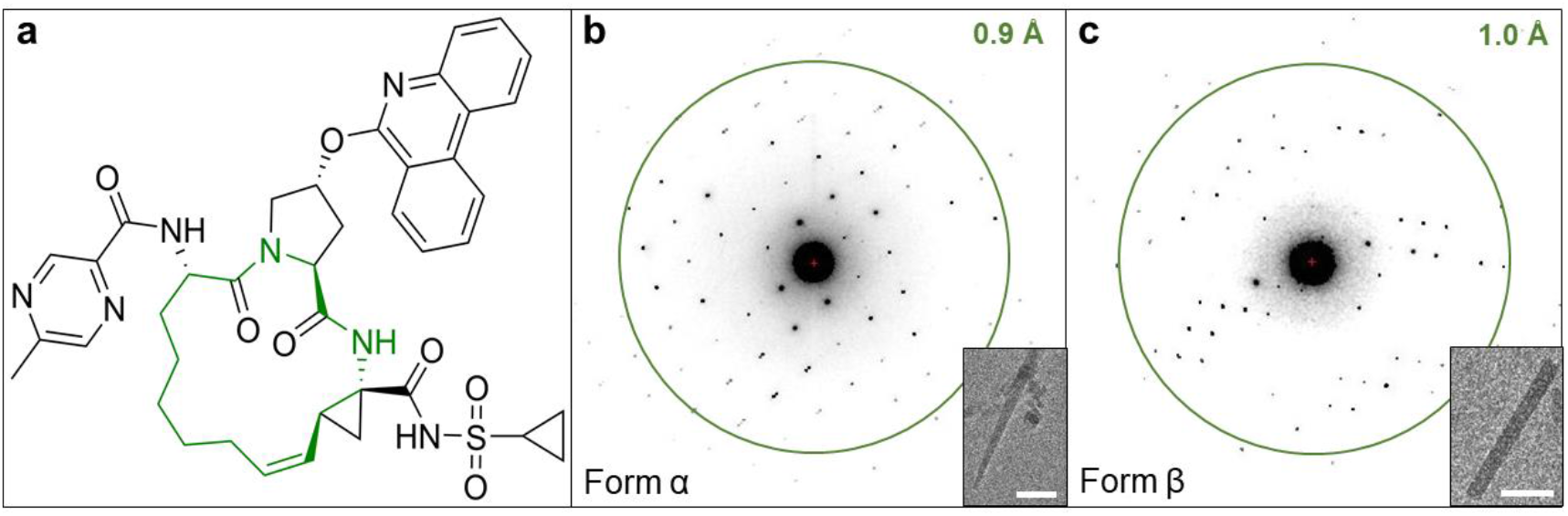
**a**: Chemical structure of paritaprevir with the macrocyclic core highlighted in green. **b:** Representative crystal image of needle-like paritaprevir form α and MicroED pattern. **c:** Representative crystal image of rod-like paritaprevir form β and MicroED pattern. Scale bars corresponds to 2 µm.

In this work, we report two polymorphic crystal forms of paritaprevir obtained from the same powder preparation and experiment, without any crystallization assay, by high-throughput MicroED using SerialEM. The new crystal structure, herein referred to as paritaprevir form β, shows a different conformation and unique packing pattern, and hence reveals a conformational polymorphism from the same sample. Molecular docking experiments shows that only form β can bind to the active site pocket of HCV NS3/4A protease and interact with the catalytic triad, whereas form α doesn’t fit into the pocket. Hence, form β is a potential target bound conformation.

### Results and discussion

#### MicroED data collection and data processing

The MicroED sample preparation and grid screening for paritaprevir was performed according to previously described protocols. [15, 24] Two major crystal forms were identified from the same grid preparation using low-magnification whole-grid atlases. Paritaprevir form α yielded needle-like microcrystals with dimensions of 4–10 µm in lengths and 0.1– 0.5 µm in widths (Figure 1b). Initial data collection confirmed that the space group and unit cell dimensions of form α match our recently published MicroED structure of paritaprevir. [15] Paritaprevir form β produced rod-like microcrystals (Figure 1c), typically 2–5 µm in lengths and 0.2–0.4 µm in widths. Using automation, MicroED data were recorded on a Thermo-Fisher Falcon III detector at an electron dose rate of 0.01 e-/Å^2^/s and 1 second exposure per frame as the sample stage was continuously rotated from -30° to +30° at 1° per second. The data were processed in XDS following previously published procedures. [24] The *ab initio* structure of form α was solved in the orthorhombic space group P2_1_2_1_2_1_ (a = 5.09 Å, b = 15.61 Å, c = 50.78 Å, and α = β = γ = 90°) from a merged dataset with an overall completeness of 89%, and refined at 0.85 Å to an R_1_ value of 0.1472 (Table S1, Figure S1). The *ab initio* structure of form β was solved in the same orthorhombic space group P2_1_2_1_2_1_ but with significantly different unit cell size (a = 10.56 Å, b = 12.32 Å, c = 31.73 Å, and α = β = γ = 90°) from a merged dataset with an overall completeness of 98.4%, and refined at 0.95 Å to an R_1_ value of 0.1347 (Table S1, Figure S3).

#### Structure analysis

Paritaprevir form α adopts an open conformation with one intramolecular hydrogen bond between the amide nitrogen on the macrocyclic core and the cyclopropylsulfonamide moiety (2.2 Å, Figure 2a and c). The packing analysis reveals that intermolecular hydrogen bonds between the amide carbonyls and amide nitrogens on the macrocyclic cores generates chain motifs along the crystallographic a axis. Further, the phenanthridine rings form intermolecular π-π interactions (Figure 2a and S2). The packing of form α gives rise to the formation of solvent accessible channels which form along the crystallographic a axis (Figure 2a). Paritaprevir form β adopts another open conformation with an intramolecular hydrogen bond between the amide carbonyl on the macrocyclic core and amide nitrogen on the cyclopropylsulfonamide moiety (2.0 Å, Figure 2b and c). As compared to form α, the packing of form β differs; intermolecular hydrogen bonds between the amide carbonyl on the methylpyrazinamide moiety and amide nitrogen on the macrocyclic core, as well as between amide nitrogen on the methylpyrazinamide moiety and aromatic nitrogen on the phenanthridine ring, generates chain motifs along the crystallographic a axis (Figure 2b and S4).

**Figure 2.**
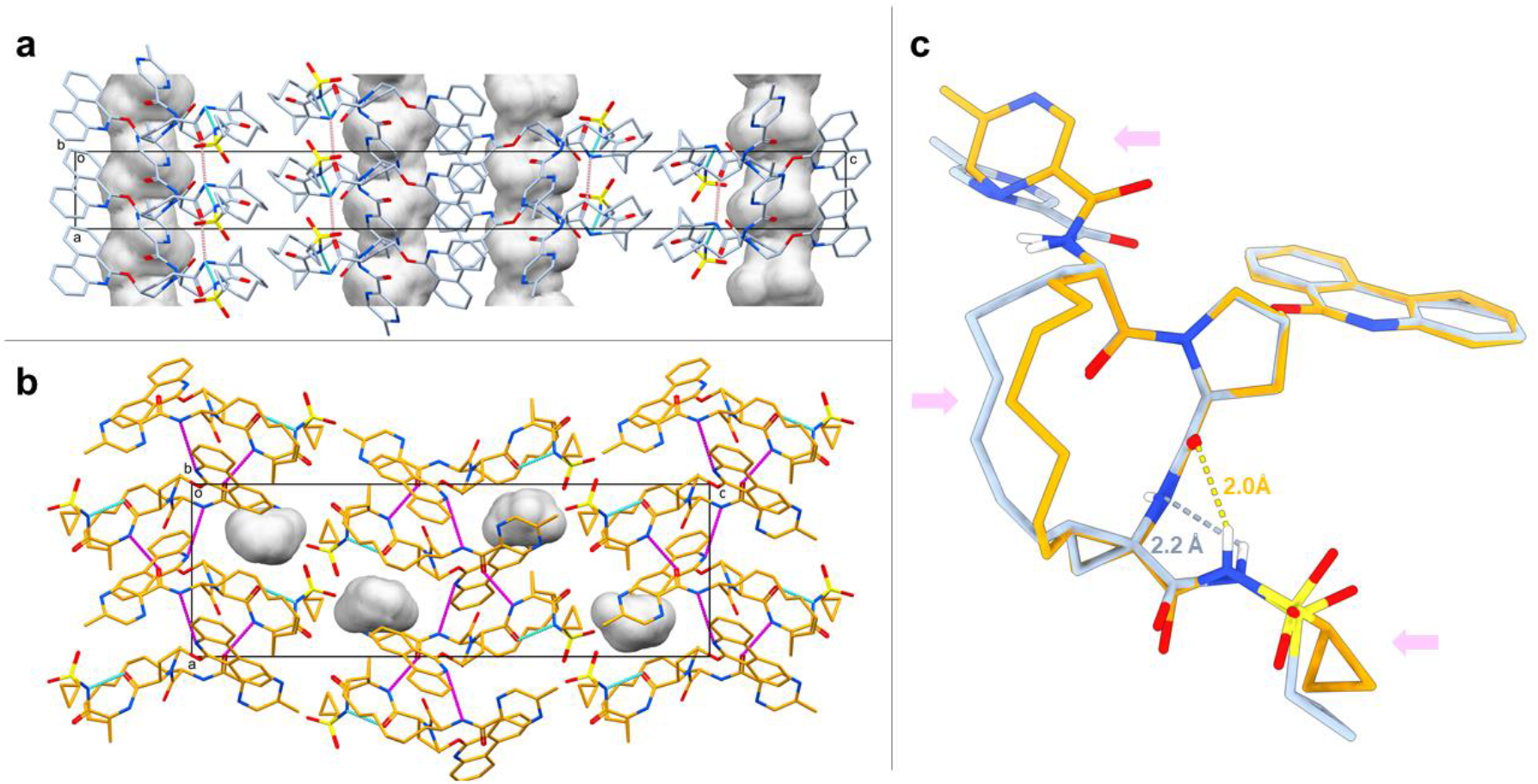
**a**: Unit cell packing of paritaprevir form α viewed along the crystallographic b axis with the unit cell box shown in black. Atom colors: C, light blue; N, blue; O, red; S, yellow. The inter- and intramolecular hydrogen bonds are shown in magenta and cyan dashed lines, respectively. The one-dimensional solvent channels are shown in gray contoured surface extending along the crystallographic a axis. All hydrogens are omitted for clarity. **b**: Unit cell packing of paritaprevir form β viewed along the crystallographic b axis with the unit cell box shown in black. Atom colors: C, orange; N, blue; O, red; S, yellow. The inter- and intramolecular hydrogen bonds are shown in magenta and blue dashed lines, respectively. The void space are shown in gray contoured surface. All hydrogens are omitted for clarity. **c**: Overlay of all heavy atoms on the aromatic rings, 5-membered ring and macrocyclic core amides between paritaprevir form α (light blue) and form β (orange), showing conformational changes in the macrocyclic core, methylpyrazinamide moiety and cyclopropylsulfonamide moiety, as indicated by the pink arrows. The intramolecular hydrogen bonds are shown in dashed lines. Non-polar hydrogens are omitted for clarity.

The observation of solvent channels in pharmaceutical crystals of molecules with chameleonic behavior has been previously described [25, 26] and can lead to reduced aqueous solubility [26] and risks in processing, manufacturing and storage of the drugs. [27] Using the default void analysis in Mercury with a probe radius of 1.2 Å, 7.6% of the unit cell for form α is concluded to consist of solvent-accessible channels extending along the crystallographic a axis (Figures 2a and S2). Paritaprevir form β does not have the large channels observed for form α. Instead, the crystal packing is significantly tighter with only 2.2% of the unit cell corresponding to discrete void space (Figure 2b and S4). The space group and unit cell dimensions of form α match those of a paritaprevir hydrate crystal form studied by powder X-ray diffraction (PXRD), [25, 28] with the unit cell axis of a and c being flipped. In the PXRD study, water molecules are located in the one-dimensional channels and interact with each other, with the sulfonamide oxygen, and with the phenanthridine nitrogen. Although the PXRD structure is not publicly accessible, form α shares a similar crystal packing with the paritaprevir hydrate form based on the same space group, unit cell dimensions, and the one-dimensional channels extending along the crystallographic axis. In contrast to the the PXRD study, paritaprevir form α is solved as an anhydrate, suggesting that it can remain the crystallinity during hydration/dehydration [28] and hence risk trapping water molecules upon storage. Taken together with the observed strong intermolecular interactions, form α is therefore assumed to have unfavorable properties for storage, solubility and dissolution rate in water, and thus is not a good candidate for formulation and process development. [25]

Although both form α and form β adopt open conformations, the structures are different with an rmsd value of 0.83 Å comparing all heteroatoms (Figure 2c). The most significant conformational changes includes the alkyl chain motifs of macrocyclic core, with a torsion angle difference over 90°, the different rotamers of the methylpyrazinamide substituents, and the cyclopropylsulfonamide moieties with slightly different orientations and significant change in torsion, resulting in different intramolecular hydrogen bond locations (Figure 2c). The distances between the centroid of the cyclopropylsulfonamide groups and olefin groups on the macrocyclic core are over 5 Å in both forms, suggesting that both forms have good chemical stability since oxidative reactions can occur if the two groups are in close proximity. [25] The calculated three dimensional polar surface area (3D-PSA) are 194.52 Å^2^ and 195.92 Å^2^ for form α and form β, respectively. This compares well to the 3D-PSA value of a simulated target bound paritaprevir conformer (186.7 Å^2^). [25] In order to understand the biological importance of both conformers, molecular docking was applied to the protein target.

#### Molecular docking

Molecular docking experiments of paritaprevir conformers α and β into HCV NS3/4A protease were performed using Glide [29] and the crystal structure of simeprevir-bound genotype 1b HCV NS3/4A (PDB ID: 3KEE [30]) as the receptor, in which the coordinates of simeprevir, water molecules, and other ligands were removed and hydrogens were added to all protein residues. This structure was selected because simeprevir (Figure S5) and paritaprevir are structurally similar and both target the HCV NS3/4A protease. Conformer α docked outside of the active site of the protease (result not shown), with a docking score of -4.2 kcal/mol, and conformer β docked into the active site pocket with a docking score of -11.3 kcal/mol (Figure 3a and b). [31-33] Hence, conformer β is more thermodynamically favored for target binding. The simeprevir-bound crystal structure revealed the antiviral to disrupt the catalytic triad of the HCV NS3/4A protease (H57, D81 and S139) by forming hydrogen bonds to side chains S139 and H57 (Figure S5). Additional hydrogen bonds are formed between simeprevir and the backbones of G137, R155 and A157, and between simeprevir and the side chain of K136, where the latter serves as a clamp to close the active site pocket (Figure S5). The structure-activity relationship study of paritaprevir and the HCV protease revealed the macrocyclic core, the cyclopropylsulfonamide moiety and the phenanthridine ring to be of high importance for the inhibitory activity. [13] The macrocyclic core interacts extensively with the pocket, similar to simeprevir, via several hydrophobic interactions as well as via hydrogen bonds to the side chain of K136, and to the backbones of R155 and A157 (Figure 3b and Table S2). The cyclopropylsulfonamide has long been incorporated into other drugs for HCV NS3/4A protease target. [13] In our model paritaprevir interacts with the catalytic triad through hydrogen bonds between the cyclopropylsulfonamide and the side chains of H57 and S139 (Figure 3b and Table S2). In addition, this substituent makes a hydrogen bond to the G137 backbone as well as to the K136 side chain, which is not present in the simeprevir structure. K136 form a clamp to control the opening of active site pocket, and G137 is the site of oxyanion hole. [34] The phenanthridine ring of paritaprevir shows hydrophobic interaction with the side chains of H57 and D81, where the latter is not present in the simeprevir-bound structure. The 5-membered ring linking the phenanthridine to the core shows additional hydrophobic interaction with the side chain of H57. Lastly, the methylpyrazinamide moiety interacts with the target via a hydrogen bond to the backbone of A157 (Figure 3b and Table S2). The few interactions of this substituent corroborates its moderate effect in improving potency, however it was shown to be important for metabolic stability and pharmacokinetics. [13] It can be noted that the potency of paritaprevir for the HCV genotype 1b target (EC_50_ = 0.21 nM) [13] is higher than for simeprevir (EC_50_ = 25 nM), [35] which might partly be explained by some of additional target interactions for paritaprevir suggested here.

**Figure 3.**
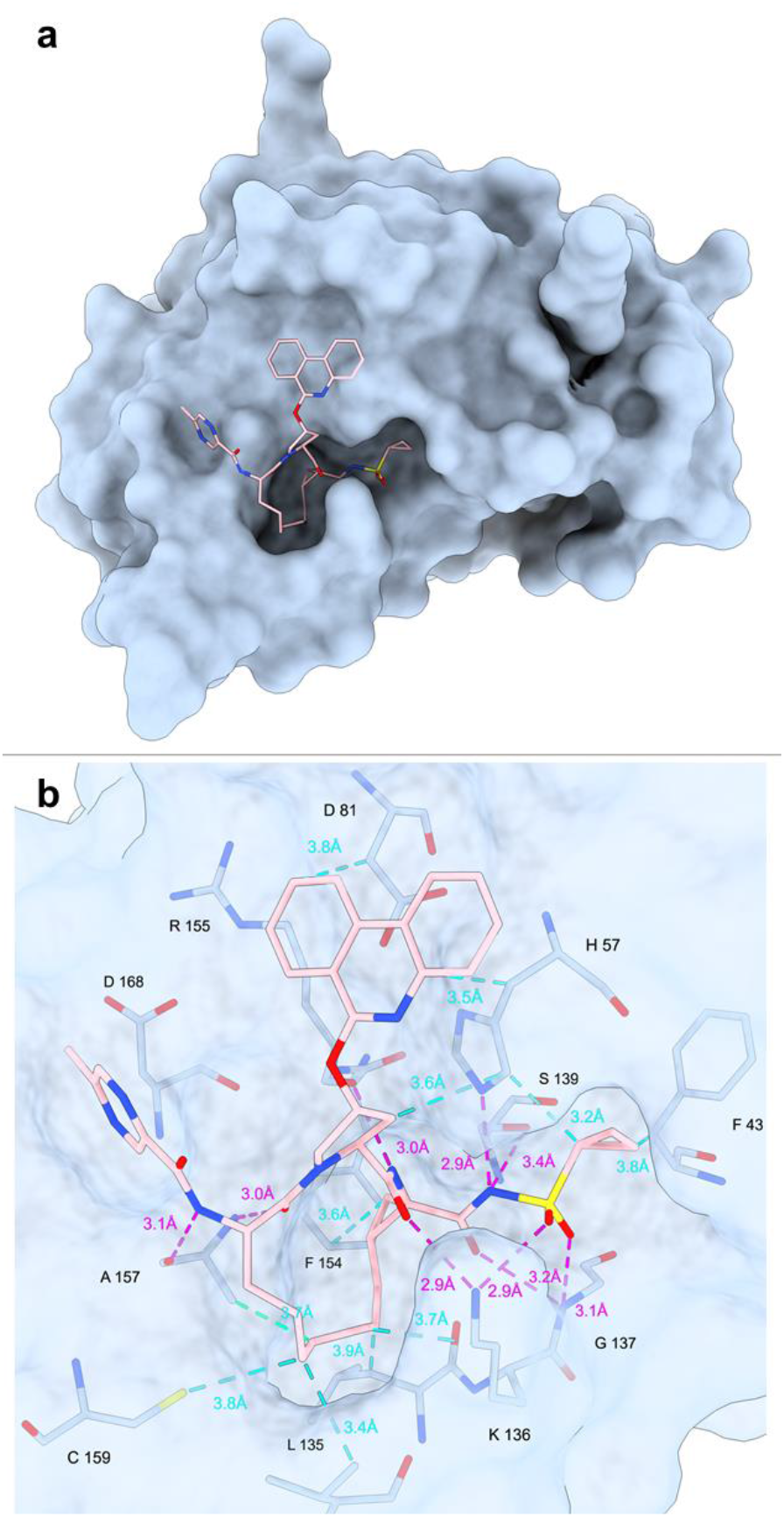
**a**: Docking of paritaprevir conformer β into the simeprevir-bound crystal structure of HCV NS3/4A protease (PDB ID: 3KEE). Atom colors: C, pink; N, blue; O, red; S, yellow. All hydrogen atoms are omitted for clarity **b:** Potential **i**nteractions between paritaprevir and the target observed from the molecular docking. The intra- and intermolecular hydrogen bonds on paritaprevir are shown as dashed lines in cyan and magenta, respectively.

## Conclusions

The development of high-throughput MicroED with automation has recently been successfully applied in drug discovery for analysis of small molecule therapeutics, [24, 36, 37] and for the structural determination of large and flexible macrocycles. [15] Herein, high-throughput MicroED is successfully used for polymorph screening from the same experiment, without crystallization assays or different sample preparations. Paritaprevir is known to have chameleonic properties and adopts open conformations in aqueous environment and closed conformations in lipophilic environment. Two of the conformations could be determined from the same CryoEM grid where one is suggested to be the target bound state. We show that high-throughput MicroED will be suitable for future polymorph screening in future drug discovery, and that MicroED structures of complex modalities in the beyond rule of 5 space can be used for precision docking to difficult-to-drug targets.

## Acknowledgement

The authors thank Johan Hattne for the setup and support of the computer cluster and its SLURM interface used for the processing. This study was supported by the National Institutes of Health P41GM136508. Portions of this research or manuscript completion were developed with funding from the Department of Defense grants MCDC-2202-002. Effort sponsored by the U.S. Government under Other Transaction number W15QKN-16-9-1002 between the MCDC, and the Government. The US Government is authorized to reproduce and distribute reprints for Governmental purposes, notwithstanding any copyright notation thereon. The views and conclusions contained herein are those of the authors and should not be interpreted as necessarily representing the official policies or endorsements, either expressed or implied, of the U.S. Government. The PAH shall flowdown these requirements to its subawardees, at all tiers. The Gonen laboratory is supported by funds from the Howard Hughes Medical Institute. E.D. thanks The Wenner-Gren Foundations for their support through the Wenner-Gren Postdoctoral Fellowship. L.W. thanks The Bengt Lundqvist Memorial Foundation, the Liljewalch foundation, and the Bergmarks foundation for fellowship support.

## Notes

### Competing Interest Statement

The authors have declared no competing interest.

