## Supplemental Information for "Polymorphic Structure Determination of the Macrocyclic Drug Paritaprevir by MicroED"

Bu G[1*], Danelius E[1,2*], Wieske L.H[3], and Gonen T[1,2,4$]

1. Department of Biological Chemistry, University of California Los Angeles, 615 Charles E.Young Drive South, Los Angeles, CA 90095, USA.
2. Howard Hughes Medical Institute, University of California Los Angeles, Los Angeles, CA 90095, USA.
3. Department of Chemistry – BMC, Uppsala University, Husargatan 3, 75237 Uppsala, Sweden.
4. Department of Physiology, University of California Los Angeles, 615 Charles E. Young Drive South, Los Angeles, CA 90095, USA.

* Denotes equal contribution

#### Experimental

**Sample preparation.** Approximately 0.5 mg paritaprevir (Invivochem) was dissolved into minimal amounts of methanol in a clean 4 mL scintillator vial, followed by solvent evaporation at room temperature for approximately 20 h. The resulting deposits were scraped from the glass wall and mixed with a pre-clipped 400-mesh TEM grid (Ted Pella) coated with continuous carbon film. Prior to grid preparation, the TEM grid was glow-discharged for 30 seconds on each side at 15 mA on the negative mode using PELCO easiGlow (Ted Pella).

**MicroED data collection.** The TEM grid was loaded into a Thermo-Fisher Talos Arctica electron microscope operating at 80 K and 200 kV (0.0251 Å wavelength). MicroED datasets were automatically collected using SerialEM following the published protocol in our recent work [1]. The whole grid atlas was acquired as low-magnification montage at a magnification of 155x. Grid squares containing microcrystals were selected by the SerialEM "Navigator", and acquired for medium-magnification montages at a magnification of 2,600x. During medium-magnification montage, the "Fine eucentricity" function in SerialEM was selected to assign the eucentric heights for each grid square in the corresponding maps. Microcrystals were picked from each medium-magnification montage within the "Navigator" window for data collection. MicroED data collection was performed in the SerialEM "Record" mode where the microscope was set for the parallel electron diffraction settings (C2 lens intensity of 45.2% inserted with an aperture size of 20, resulting in the beam size of approximately 1.5 µm in diameter). Continuous-rotation MicroED data were recorded as MRC format on a Thermo-Fisher Falcon III detector in linear mode at an electron dose rate of 0.01 e-/(Å$^2$*s)) and 1 second exposure per frame as the sample stage was tilting from -30˚ to +30˚ at 1˚ per second.

**MicroED data processing and structure determination.** The datasets were initially processed using an in-house developed python script for automatic image conversion, indexing, integration and scaling [1, 2]. The information on completeness, resolution and cell dimensions generated by our automatic script provided a guideline for which data sets could be manually processed and merged in XDS [3] in order to improve the processing and merging statistics. Reflection files were prepared using XPREP (Bruker), and the *ab initio* structures were determined by SHELXD [4], followed by refinement in SHELXL [5] using electron scattering factors. Unless specified, hydrogen atoms were located at the geometrically idealized positions and refined using riding model. MicroED structures were visualized in ChimeraX [6]. Crystal packing and RMSD were analyzed in Mercury [7]. 3D polar surface area were calculated using Qikprop (Schrodinger).

**Molecular docking.** Molecular docking was performed in Maestro (Schrodinger) using the published crystal structure of simeprevir-bound Hepatitis C virus (HCV) NS3/4A protease (PDB ID: 3KEE [8]) as the receptor. At first, the default settings of Protein Preparation

#### *SUPPLEMENTARY INFORMATION*

Wizard were employed for preparing receptor molecule for docking. Simeprevir, water and other ligand molecules were removed from the receptor. Hydrogens were added to the receptor and optimized using the PROPKA algorithm. The receptor was then energy-minimized using the OPLS4 force field. The next procedure was to generate the receptor grid for docking using a box size of 36 Å in length centered at the H57 residue of NS3 chain. The van der Waals radii of nonpolar receptor atoms were scaled at a factor of 1.0 using partial charge cutoff at 25%. Lastly, docking experiments were performed using Glide (Schrodinger) [9] in the extra precision (XP) mode. Paritaprevir conformer α and β were set as flexible. Nitrogen inversions and the macrocycle ring conformations were sampled. The van der Waals radii of nonpolar ligand atoms were scaled at a factor of 0.8 using partial charge cutoff at 30%. Conformer α was docked outside of the active site pocket, with a score of -4.2 kcal/mol. Conformer β was docked into the active site pocket, with a score of -11.3 kcal/mol. The docked model and protein-drug interactions were visualized in ChimeraX [6].

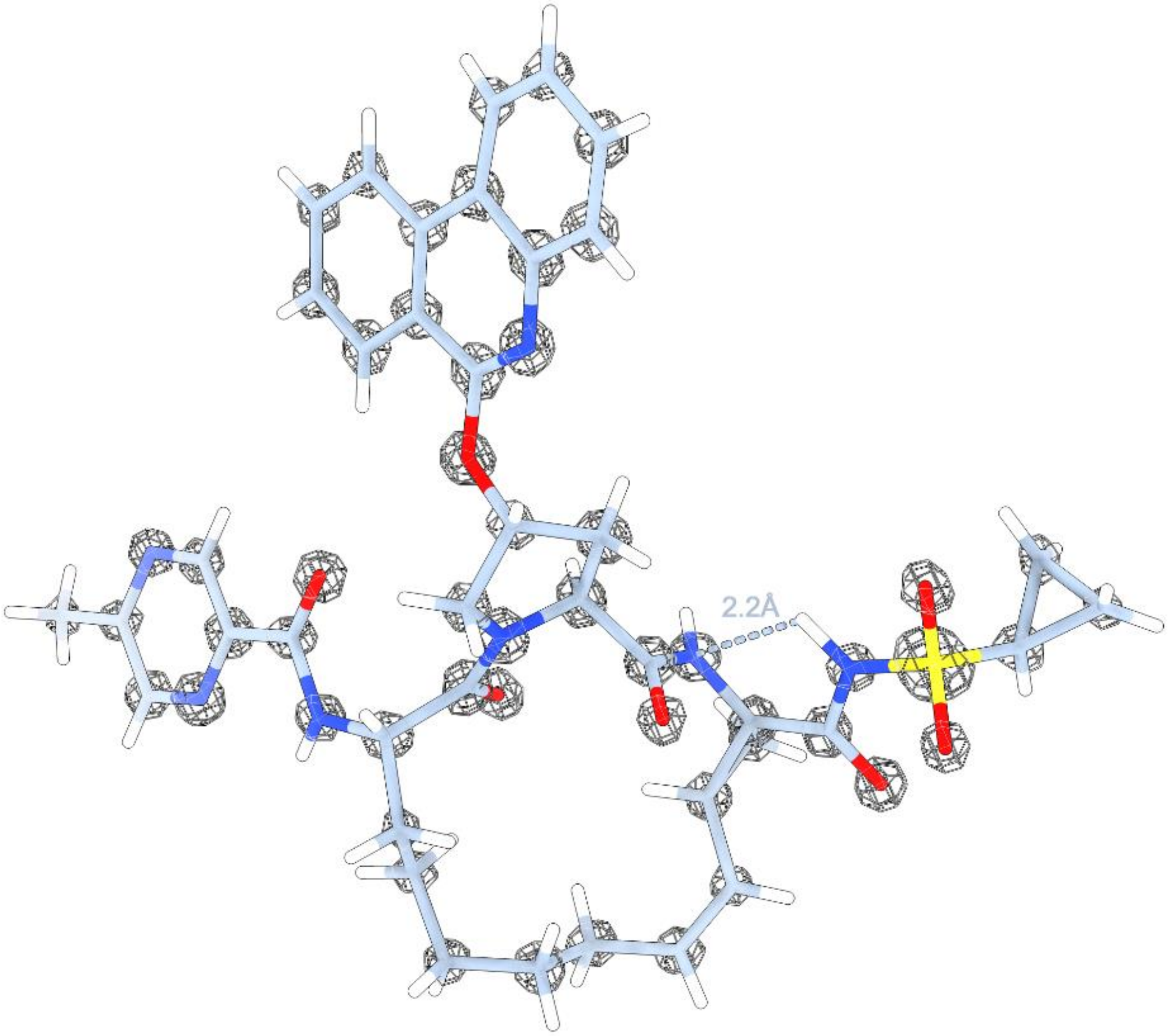


**Figure S1:** MicroED structure of paritaprevir form α with the density contoured at 1.5σ level. Atom color: C, gray; N, blue; O, red; S, yellow; H, white. Densities are shown as meshed contoured surface in light steel blue.

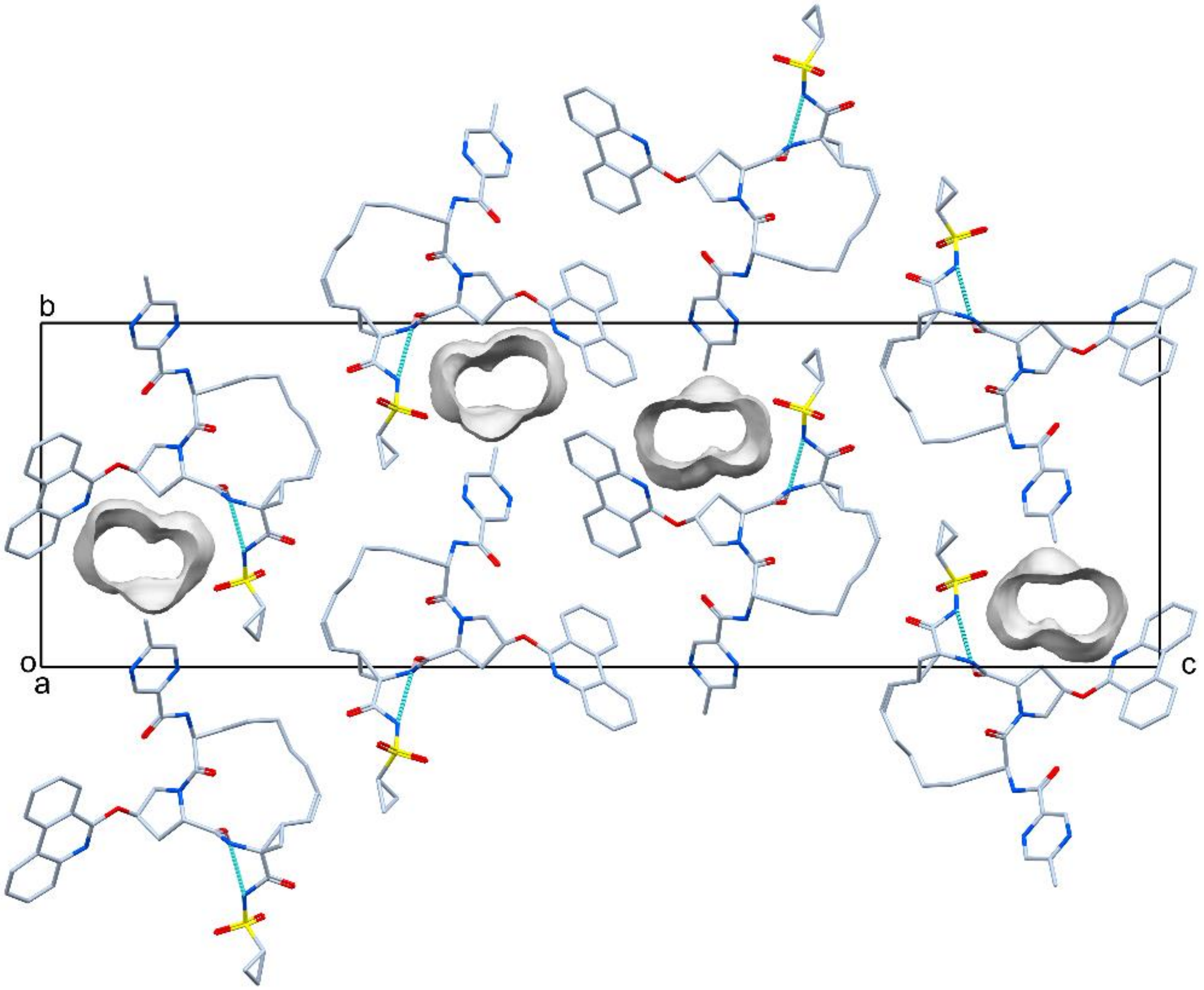


**Figure S2:** Unit cell packing of paritaprevir form α viewed along the crystallographic a axis with the unit cell box shown in black. Atom colors: C, light steel blue; N, blue; O, red; S, yellow. The inter- and intramolecular hydrogen bonds are shown in magenta and cyan dashed lines, respectively. The one-dimensional channels are shown in light gray contoured surface extending along the crystallographic a axis. All hydrogens are omitted for clarity.

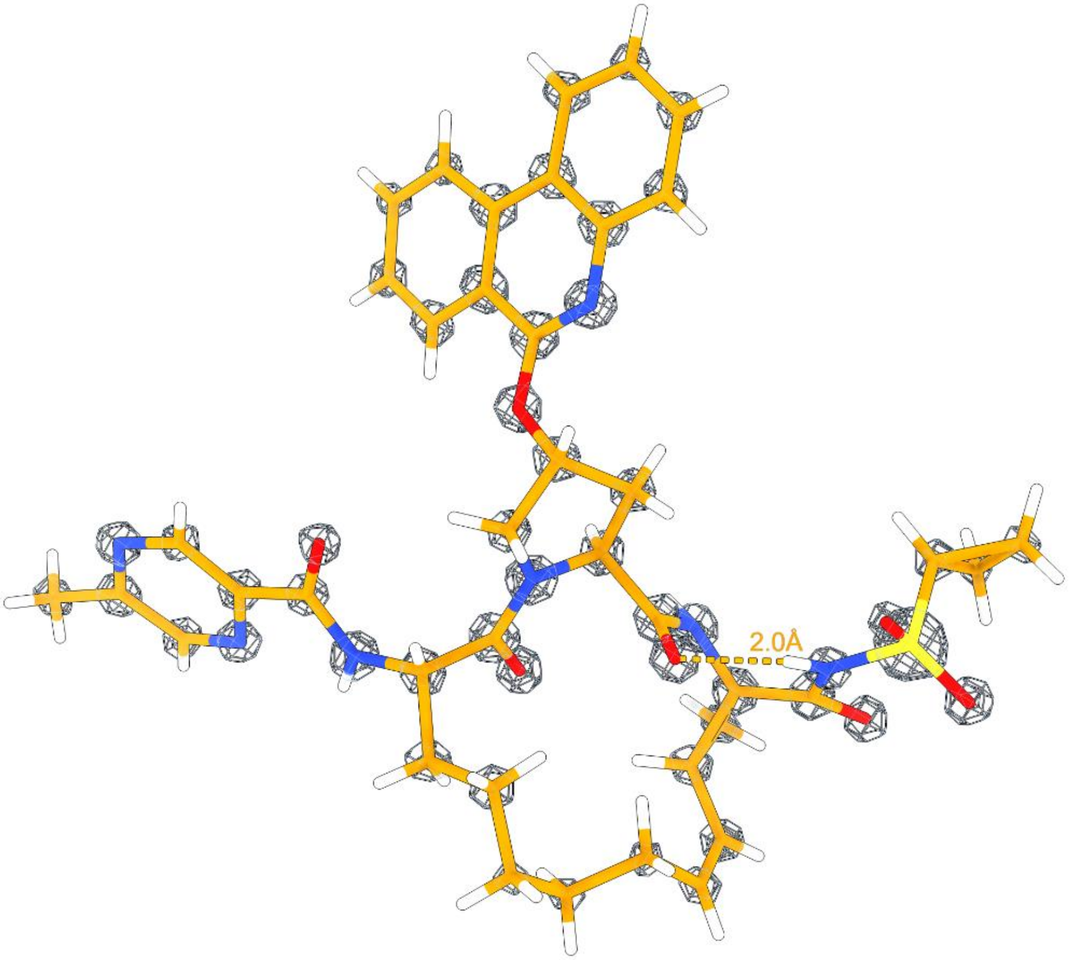


**Figure S3:** MicroED structure of paritaprevir form β with the density contoured at 1.5σ level. Atom color: C, orange; N, blue; O, red; S, yellow; H, white. Densities are shown as meshed contoured surface in light steel blue.

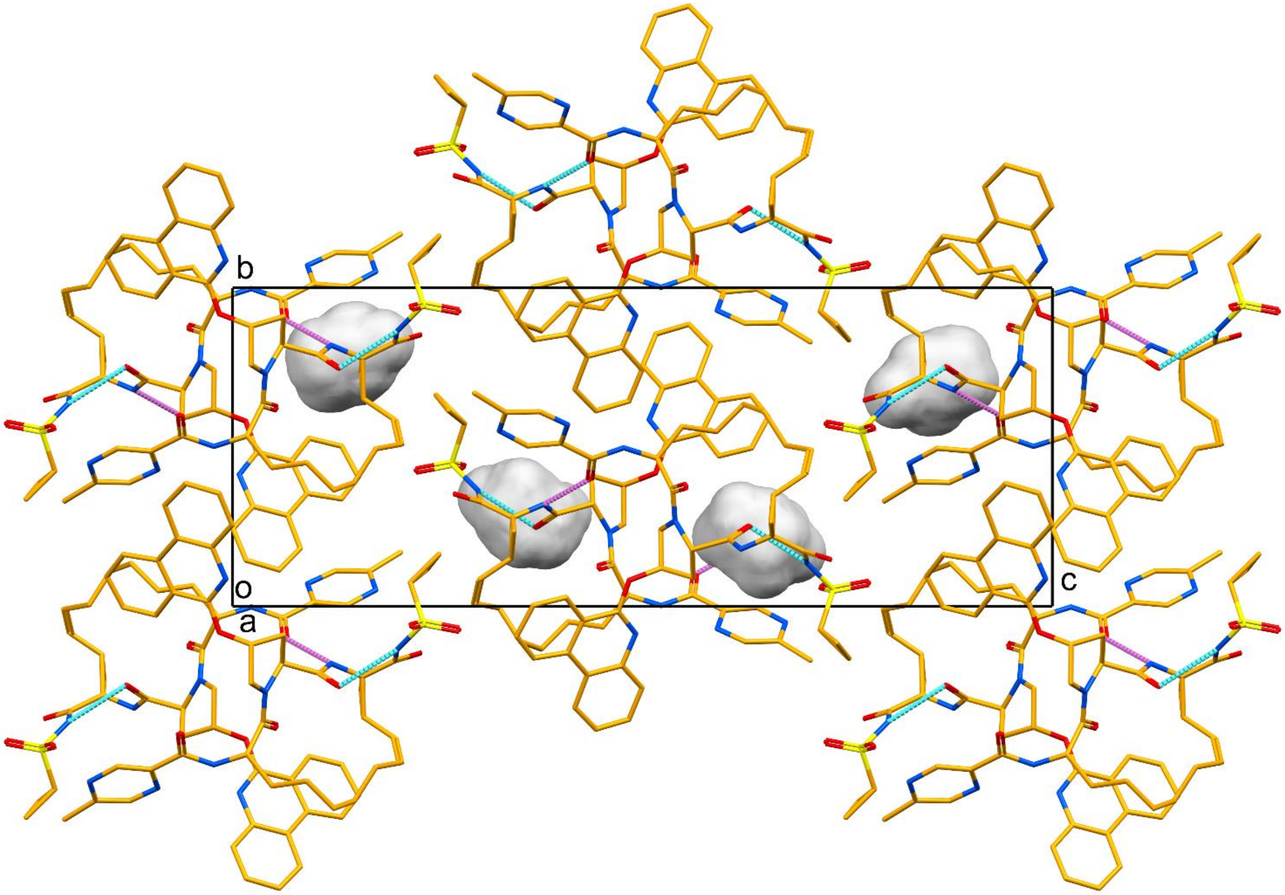


**Figure S4:** Unit cell packing of paritaprevir form β viewed along the crystallographic a axis with the unit cell box shown in black. Atom colors: C, orange; N, blue; O, red; S, yellow. The inter- and intramolecular hydrogen bonds are shown in magenta and cyan dashed lines, respectively. The voids are shown in light gray contoured surface. All hydrogens are omitted for clarity.

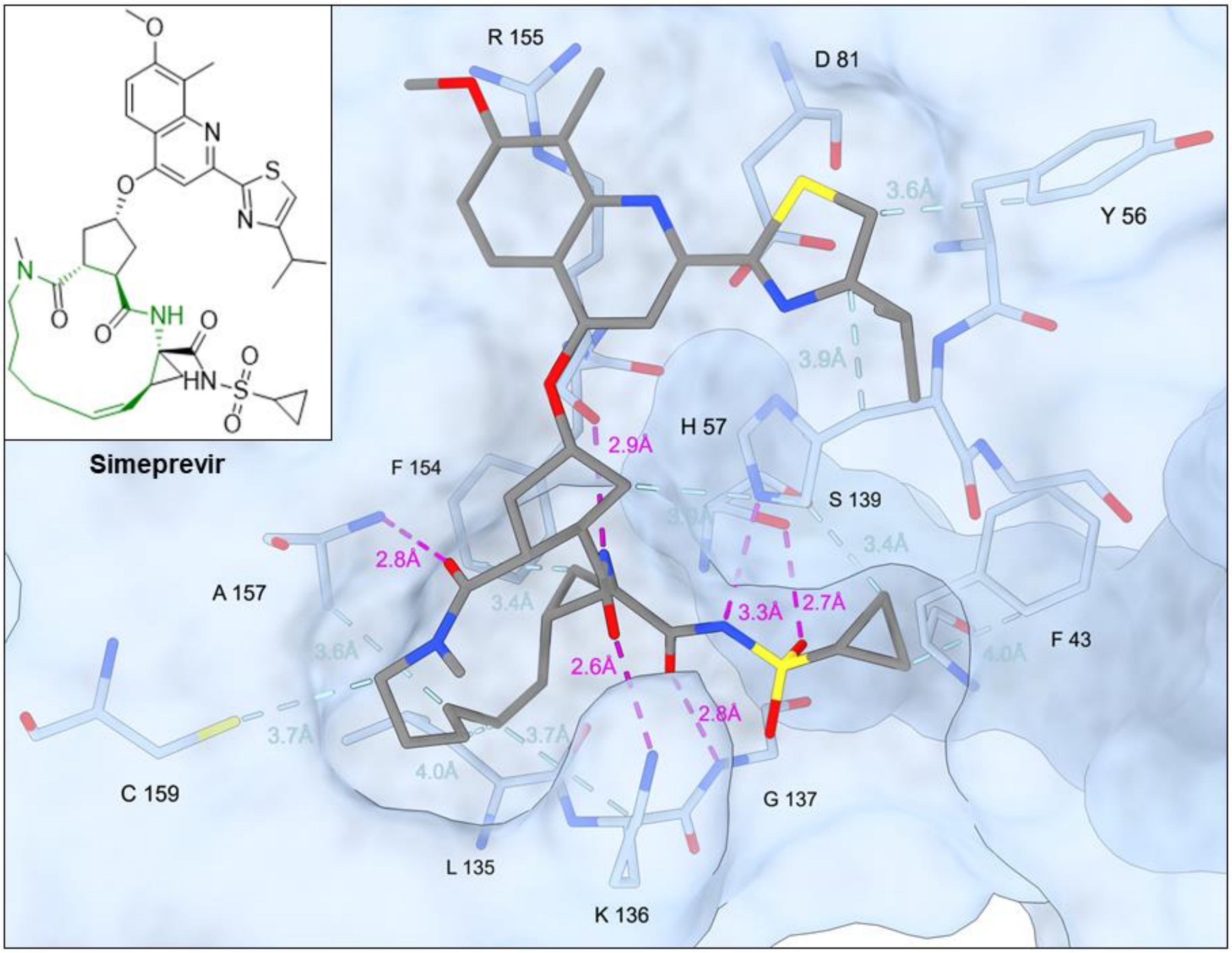


**Figure S5:** Interactions between simeprevir and HCV NS3/4A protease observed from the X-ray structure (PDB ID: 3KEE). Atom color in simeprevir: C, gray; N, blue; O, red; S, yellow. Hydrophobic interactions and hydrogen bonds are shown as light blue and magenta dashed lines, respectively. All hydrogen atoms are omitted for clarity.

### *SUPPLEMENTARY INFORMATION*

**Table S1**. Processing and structure refinement statistics of paritaprevir form α and β

| | **Form α** | **Form β** |
|---|---|---|
| Stoichiometric formula | $C_{40}H_{43}N_7O_7S$ | $C_{40}H_{43}N_7O_7S$ |
| Radiation wavelength (Å) | 0.0251 | 0.0251 |
| Temperature (K) | 80 | 80 |
| Number of crystals | 2 | 2 |
| Crystal shape | needle | rod |
| Resolution (Å) | 50.0 - 0.85 | 50.0 - 0.95 |
| Crystal system | orthorhombic | orthorhombic |
| Space group | $P2_12_12_1$ | $P2_12_12_1$ |
| Unit cell length a, b, c (Å) | 5.09, 15.61, 50.78 | 10.56, 12.32, 31.73 |
| Unit cell angle α, β, ɣ (°) | 90, 90, 90 | 90, 90, 90 |
| Z | 4 | 4 |
| Total reflections | 15,086 | 16,900 |
| Unique reflections | 3,607 | 2,851 |
| $R_{obs}$ (%) | 20.2 | 22.8 |
| $R_{meas}$ (%) | 23.2 | 24.9 |
| I/σI | 5.20 | 5.41 |
| $CC_{1/2}$ (%) | 98.9 | 98.9 |
| Completeness (%) | 89.0 | 98.4 |
| $R_1$ | 0.1472 | 0.1347 |
| $wR_2$ | 0.4080 | 0.3734 |
| GooF | 1.156 | 1.114 |
| CCDC number | TBD | TBD |

**Table S2**. Statistics of interactions between paritaprevir and HCV NS3/4A protease observed from molecular docking.

| Interactions | Receptor atom (residue) | Ligand atom (moiety) | Distance in Å |
|---|---|---|---|
| **Hydrogen bond** | NE2 (H57) | N3 (cyclopropylsulfonamide) | 2.9 |
| | NZ (K136) | O2 (macrocyclic core) | 2.9 |
| | NZ (K136) | O5 (cyclopropylsulfonamide) | 2.9 |
| | N (G137) | O4 (cyclopropylsulfonamide) | 3.1 |
| | N (G137) | O3 (cyclopropylsulfonamide) | 3.2 |
| | OG (S139) | N3 (cyclopropylsulfonamide) | 3.4 |
| | O (R155) | N2 (macrocyclic core) | 3.0 |
| | N (A157) | O1 (macrocyclic core) | 3.0 |
| | O (A157) | N4 (methylpyrazinamide) | 3.1 |
| **Hydrophobic interaction** | CB (F43) | C26 (cyclopropylsulfonamide) | 3.8 |
| | CD2 (H57) | C25 (cyclopropylsulfonamide) | 3.2 |
| | CB (H57) | C40 (phenanthridine) | 3.5 |
| | CD2 (H57) | C3 (5-membered core) | 3.6 |
| | CB (D81) | C32 (phenanthridine) | 3.8 |
| | CG1 (V132) | C13 (macrocyclic core) | 3.4 |
| | CB (L135) | C9 (macrocyclic core) | 3.9 |
| | CD (K136) | C11 (macrocyclic core) | 3.7 |
| | CZ (F154) | C7 (macrocyclic core) | 3.6 |
| | CB (A157) | C12 (macrocyclic core) | 3.7 |
| | SG (C159) | C13 (macrocyclic core) | 3.8 |
